# Seasonal climate variations promote bacterial α-diversity in soil

**DOI:** 10.1101/2020.08.04.234278

**Authors:** Xin-Feng Zhao, Wen-Sheng Shu, Yi-Qi Hao

**Affiliations:** School of Life Sciences, South China Normal University, Guangzhou 510631, China

**Keywords:** climate seasonality, coexistence, diversity, fluctuation-dependent mechanisms, soil bacteria

## Abstract

Ecological theory suggests that temporal environmental fluctuations can contribute greatly to diversity maintenance. Given bacteria’s short generation time and rapid responses to environmental change, seasonal climate fluctuations are very likely to play an important role in maintaining the extremely high α-diversity of soil bacterial community, which has been unfortunately neglected in previous studies. Here, with in-depth analyses of two previously published soil bacterial datasets at global scale, we found that soil bacterial α-diversity was positively correlated with both seasonal variations of temperature and precipitation. Furthermore, piecewise structural equation models showed that seasonal variations of temperature or precipitation directly promoted soil bacterial α-diversity in each dataset. However, it is noteworthy that the importance of seasonal climate variations might be underestimated in the above analyses, due to the high level of environmental variations irrelevant to climate seasonality among sampling sites and the lack of sampling across seasons. Supplementary analyses of a previously published wheat cropland dataset with samples collected in both summer and winter across North China Plain was conducted. Similarly, we found that bacterial α-diversity was positively correlated with seasonal climate variations, but much stronger, though the range of seasonal climate variations was much smaller compared to the two global datasets. Collectively, these findings implied that fluctuation-dependent mechanisms of diversity maintenance presumably operate in soil bacterial communities. Based on existing evidence, we speculated that the storage effect may be the main mechanism responsible for diversity maintenance in soil bacterial community, but rigorous experimental tests are needed in the future.

## Main text

The extraordinarily high α-diversity of soil bacterial communities has fascinated and puzzled microbial ecologists [1, 2]. In contrast to the considerable advances in characterizing soil bacterial biogeographic patterns across spatial and environmental gradients (e.g. [3, 4]), little attempt has been made to infer the underlying mechanisms of diversity maintenance. The high spatial heterogeneity of soil was generally thought as an important factor in maintaining bacterial α-diversity [5]; surprisingly, temporal environmental fluctuations promoting diversity maintenance firstly suggested by Hutchinson was rarely mentioned, which is one of the most influential ideas in community ecology [6, 7]. The fluctuation-dependent coexistence mechanisms are particularly important for micro-organisms due to their short generation time and rapid responses to environmental changes [8, 9]. Temporal changes in temperature and precipitation can significantly alter soil physiochemical and nutritional status, which jointly enhance temporal environmental fluctuations in soil, and drive the seasonal species turnover in bacterial community [10]. However, the majority of biogeographic studies only chose the mean annual temperature (MAT) and annual precipitation (AP) as climatic predictors of soil bacterial diversity and composition, whereas seasonal climate variations were unfortunately neglected. Here we tested the prediction that seasonal climate variations promote soil bacterial α-diversity derived from the fluctuation-dependent coexistence theory, with two global datasets covering wide geographic and climate gradients: global topsoil microbiome (hereafter “topsoil data”) [3] and global soil atlas (hereafter “atlas data”) [11], and one regional dataset of wheat cropland soil across North China Plain (hereafter “farmland data”) [12]. Details of data acquisition, processing, and statistical analyses were provided in Supplementary Methods.

The climate data were obtained from WorldClim2 database [13], whereby the standard deviation of monthly mean temperature and the coefficient of variation of monthly precipitation were used to represent temperature seasonality (TS) and precipitation seasonality (PS), respectively. Soil bacterial richness was measured by the number of observed exact sequence variants (ESVs). As predicted, soil bacterial richness significantly increased with TS and PS in both global datasets (Fig. 1). Piecewise structural equation models [14] were then fitted to address the direct and indirect effects of geographic (absolute latitude), climatic (MAT, AP, TS, PS) variables and soil physiochemical properties (pH and C/N ratio, the most important two soil physiochemical factors in determining bacterial α-diversity and composition) on soil bacterial richness. The modeling results demonstrated significant direct effect of PS (Fig. 2a, Table S3; standardized coefficient *β* = 0.195, *p* = 0.0019) in promoting soil bacterial richness in topsoil data, and TS (Fig. 2b, Table S5; standardized coefficient *β* = 0.191, *p* = 0.0006) in atlas data after controlling for the effects of MAT, AP, soil pH and C/N ratio. Given that richness estimation is sensitive to detection of rare species, above analyses were repeated with Shannon diversity, which showed similar results as richness (Fig. S3, S4, Table S4, S6).

**Fig. 1.**
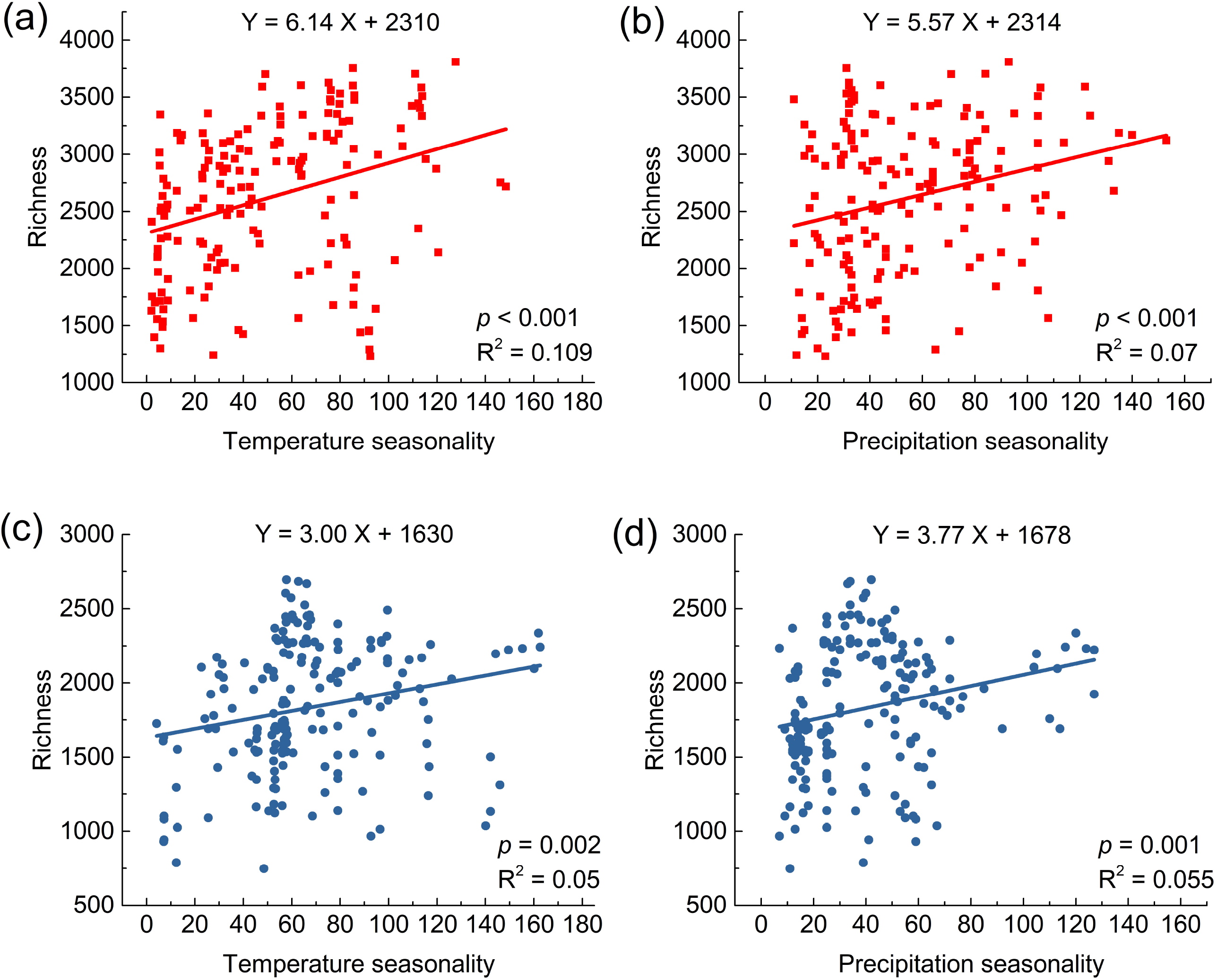
The relationships between soil bacterial richness (the number of observed ESVs) and temperature (a, c) or precipitation (b, d) seasonality in two global datasets (topsoil data: red squares, atlas data: blue circles). Significant relationships are depicted by regression lines

**Fig. 2.**
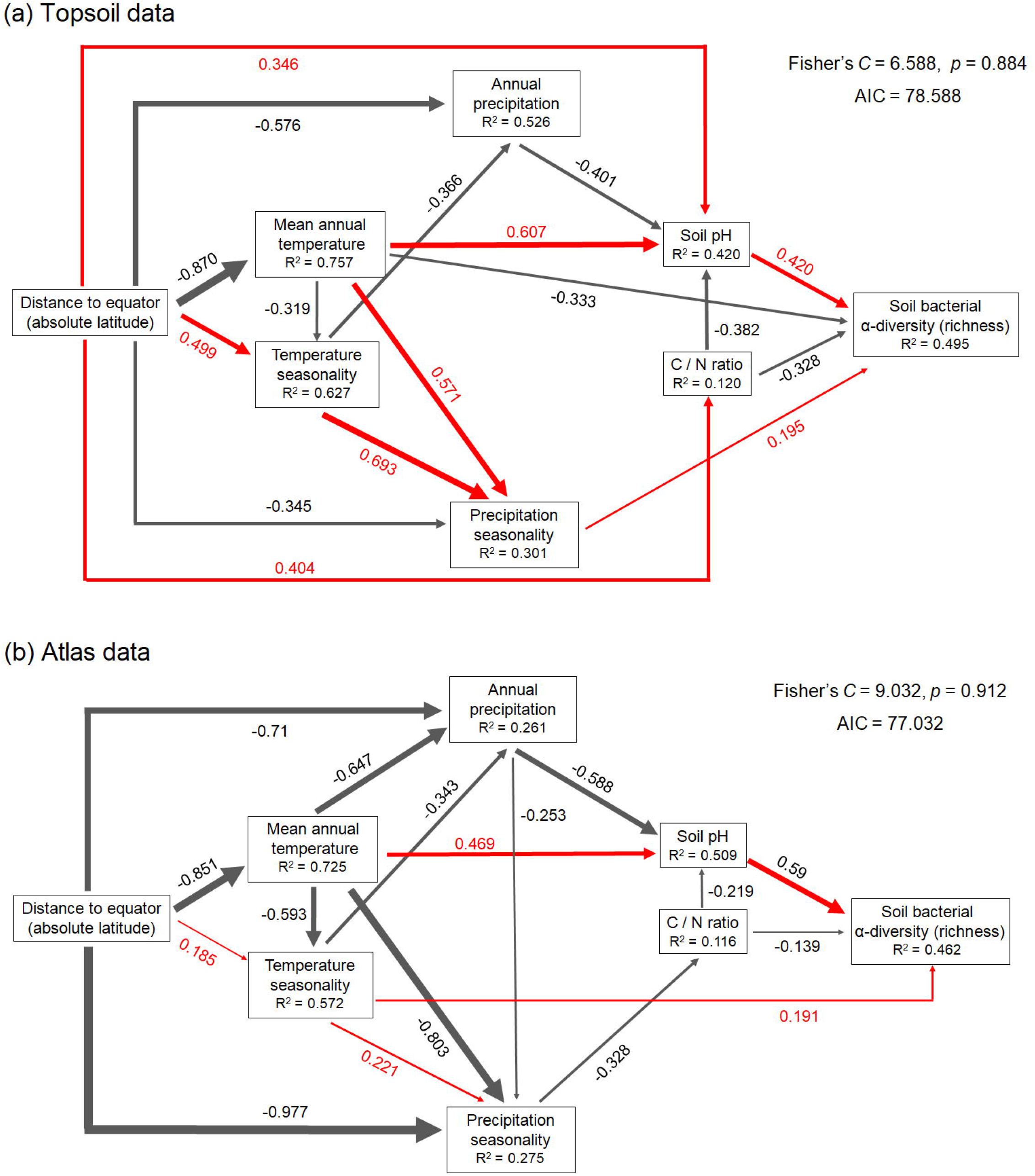
Structural equation models (piecewise SEM) to address direct and indirect effects of geographic (absolute latitude), climatic (mean annual temperature, annual precipitation, temperature seasonality, precipitation seasonality) variables, and soil physiochemical properties (pH, C/N ratio) on soil bacterial richness for topsoil data (169 soil samples) (a) and atlas data (186 soil samples) (b). Red arrows represent positive paths, and black arrows represent negative paths; only significant paths are shown (*p* < 0.05). Standardized effect sizes of path coefficients are reported and indicated by path thickness. R^2^ for component models are given in the boxes of corresponding response variables. Shipley’s test of directed separation: Fisher’s *C* statistic (if *p* > 0.05, then there are no missing associations and the model reproduces the data well) and AIC are used to evaluate the overall fit of the model. Details of model evaluation, simplification and coefficients estimation are provided in supplementary methods and Table S3, S5

Of note, the above two global datasets were consisted of soil samples from various types of ecosystem, where each location was represented by one sample [3, 11]. The high level of environmental variations between samples and lack of sampling across seasons may result in underestimating the effects of climate seasonality on maintaining soil bacterial α-diversity. As a supplement, we analyzed farmland data with soil samples collected in both summer and winter across North China Plain [12]. We found that soil bacterial α-diversity in summer was positively correlated with the collective α-diversity of two seasons (Fig. 3a, Fig. S5a). Besides, both the summer α-diversity and whole year α-diversity of soil bacterial communities were positively correlated with TS and PS (Fig. 3b, c, Fig. S5b, c). Such results enhanced our confidence in the global analyses with cross-sectional data. Despite the narrow range of climate seasonality in farmland data compared to the two global datasets, climate seasonality explained a much larger portion of variations of the whole year soil bacterial α-diversity within one type of ecosystem (Fig. 3b, c; Fig. S5b, c), where soil samples shared similar properties. This finding suggested that the importance of seasonal climate variations in maintaining soil bacterial α-diversity was probably underestimated in the global analyses. Unfortunately, the scarcity of studies incorporating seasonal variations in soil bacterial α-diversity across climate gradients impeded further meta-analysis.

**Fig. 3.**
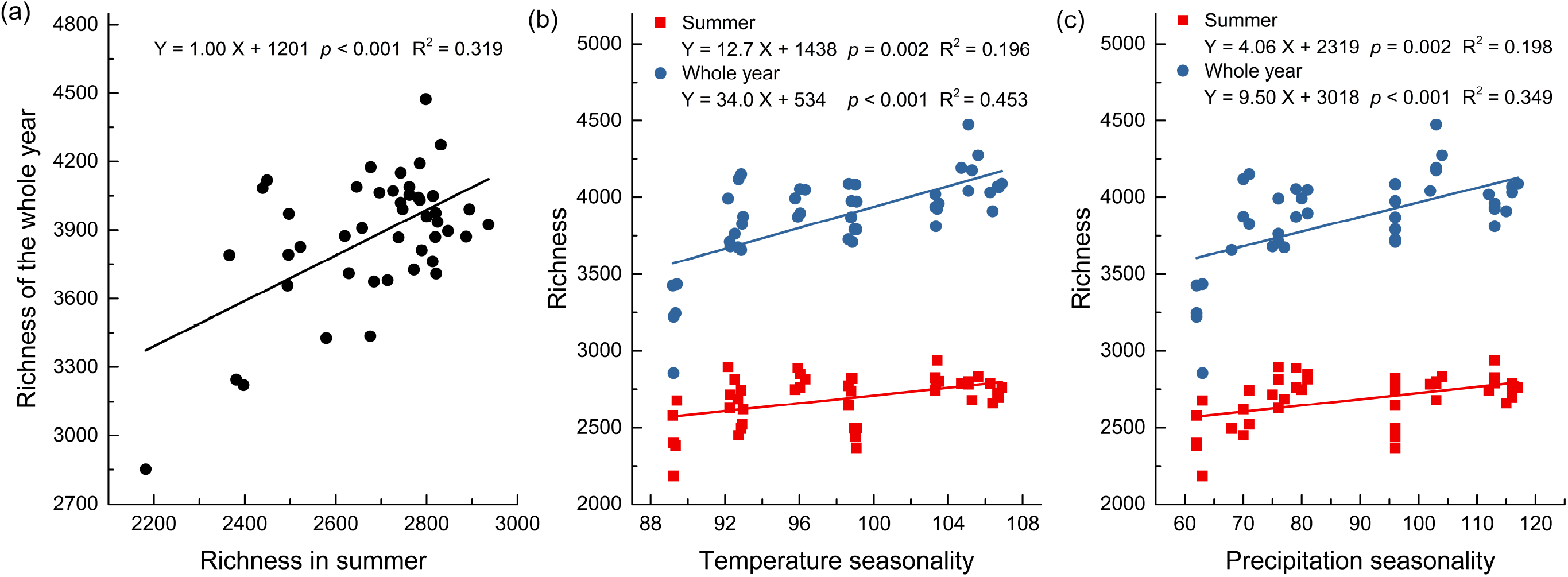
The relationship between soil bacterial richness in summer and of the whole year (a), and the relationships between soil bacterial richness (summer: red squares; whole year: blue circles) and temperature (b) or precipitation (c) seasonality. In total of 90 soil samples were collected from 45 soil plots (geographic distances up to 873 km) across North China Plain in winter and the following summer. Richness of the whole year is calculated by merging community compositions in summer and winter for each plot. Significant relationships are depicted by regression lines

Theoretically, there are two classes of fluctuation-dependent coexistence mechanisms: the storage effect and relative nonlinearity of competition [6, 9]. The storage effect mediates coexistence via temporal niche partitioning [6], and its operation requires that (1) species differ in their responses to environments, which results in (2) the relative strength of interspecific and intraspecific competition varying with fluctuating environments; and that (3) there are mechanisms buffering population from extinction under unfavorable conditions [6], such as a variety of physiological stress-resistance and dormancy mechanisms of soil bacteria [15]. There has been much experimental evidence that soil bacteria show diverse responses to moisture and temperature gradients. For example, a collection of soil bacterial isolates exhibited a wide range of responses to moisture gradient, and the derived niche parameters suggested a potential of coexistence via partitioning the moisture niche axis [16]. In addition to dry- and wet-adapted bacteria, stress-tolerant and opportunistic strategies of soil bacteria in response to drought or raining events were also discovered, as well as sensitive taxa [15, 17]. Likewise, both warm- and cold-responsive taxa were identified [18], and soil bacterial communities can rapidly diverge under contrasting experimental temperature treatments [19]. Besides evidence strongly supporting differential responses to moisture and temperature (requirement 1) discussed above, repeated reports of seasonal turnover in soil bacterial community structure (e.g. [10]) imply altered competitive intensities among soil bacteria under fluctuating environments (requirement 2); although distinguishing whether it is due to the fluctuating temperature and moisture *per se*, and/or the accompanying seasonal changes in availability of various resources is difficult. So far, rigorous tests of the storage effect have been restricted to simplistic microcosms [20, 21]; although being presumably important, its role in maintaining soil bacterial α-diversity in natural system remains unexamined yet.

By contrast, relative nonlinearity of competition might not be the major mechanism responsible for the promoting effect of seasonal climate variations on the soil bacterial α-diversity. Coexistence via relative nonlinearity of competition relies on that the per capita growth rates of species are different nonlinear functions of limiting resource [6, 8, 22]. Such nonlinearity can result in the fluctuation of limiting resource, and the stable coexistence can achieve when each species is disadvantaged relative to the others in the fluctuation environment caused by itself, but not necessarily requires fluctuation of external conditions [6, 8, 22]. On the other hand, previous modeling investigations emphasizing the importance of relative nonlinearity of competition merely focused on microorganisms in homogenous aquatic systems [8, 9], and their implications cannot be directly generalized to the soil environment with high spatial heterogeneity, where resources are not simultaneously accessible to all species.

In conclusion, our findings supported the idea that seasonal temperature and precipitation variations promote soil bacterial α-diversity via fluctuation-dependent mechanisms of diversity maintenance. Our study highlighted the importance of considering both the average values and temporal variations of abiotic environmental factors to understand the soil bacterial community composition. Further filed investigations of spatial-temporal turnover of soil bacterial diversities and experimental studies designed to investigate specific coexistence mechanisms will shed new light on the diversity maintenance of soil bacterial communities in the future.

## Supporting information

Supplementary methods and results

## Acknowledgements

We thank Mohammad Bahram for generously sharing the metadata with us.

