## Supplementary methods and results for "Seasonal climate variations promote bacterial α-diversity in soil"

Postal address: No.55 Zhongshan Avenue West, Tianhe District, Guangzhou 510631, China

### Supplementary Methods

#### Data acquisition and processing

We tested the hypothesis that seasonal temperature and precipitation variations promote soil bacterial α-diversity with two previously published soil bacterial 16S rRNA gene sequencing data at global scale: global topsoil microbiome [1] (hereafter referred to as “topsoil data”) and global soil atlas [2] (hereafter referred to as “atlas data”) and one regional dataset of wheat cropland soil across North China Plain [3] (hereafter referred to as “farmland data”). The two global datasets were chosen because they have represented the largest sampling scale across globe so far, capturing a substantial range of geographic, climatic variations and soil types; and each has consistent and standardized protocols for sampling, soil properties analyzing, DNA purification and sequencing [1, 2]. However, they were consisted of soil samples collected from various ecosystem types with high level of environmental heterogeneity irrelevant to seasonal climate variations, and no temporal sampling across seasons. Hence, supplementary analyses were then performed with farmland data which were consisted of 90 wheat cropland soil samples collected in both winter and summer from 45 plots across North China Plain (geographic distance spanned up to 878 km) [3].

Raw reads of topsoil data were downloaded from the European Bioinformatics Institute Sequence Read Archive database, under accession numbers PRJEB19856 (ERP021922); and the metadata including geographic information and soil physiochemical properties were kindly provided by the corresponding author. Raw reads and metadata of atlas data are available in Figshare (https://figshare.com/s/82a2d3f5d38ace925492; DOI: 10.6084/m9.figshare.5611321). Raw reads of farmland data were obtained from the National Center of Biotechnology Information (NCBI)

Sequence Read Archive, under accession number PRJNA508409. Climate data including mean annual temperature, annual precipitation, temperature seasonality, and precipitation seasonality at a spatial resolution of 0.86 $km^2$ were obtained from the WorldClim2 database (https://www.worldclim.org/data/monthlywth.html) [4]. The standard deviation of monthly mean temperature and the coefficient of variation of monthly precipitation were used to represent temperature and precipitation seasonality, respectively. Climate information for four samples in topsoil data were unavailable, and they were removed from the subsequent analyses.

The raw sequencing data were processed using QIIME2 2019-07 [5]. The paired-end reads were firstly assembled, and the *qiime quality-filter q-score-joined* was then applied to filter and trim the paired joined sequences based on quality scores and the presence of ambiguous nucleotide bases with the following parameters: $r$ = 3, $q$ = 4, $p$ = 0.75, and $n$ = 0 [6]. The filtered sequences were denoised using the *qiime deblur denoise-16S* method [7] with default parameters except for the trimming length parameter. Sequences of the farmland data were trimmed to 250 bp based on their high qualities. Whereas sequences of the topsoil and atlas data were trimmed to 150 bp based on the quality of sequences, considering that 150 bp sequences reveal sufficient detailed diversity patterns of microbial community composition [8] and meanwhile allow more samples with more than 10 000 quality controlled reads included in the analyses. Deblur algorithm identified exact sequence variants (ESVs) that differed by a single base instead of clustering OTUs at 97% similarity, and ESVs with less than 10 reads across all samples in each dataset were removed to minimize the impacts of sequencing errors. The *de novo* chimeras filtering was achieved with VSEARCH (parameters: dn = 0.000001, xn = 1000, minh = 10000000, mindiffs = 5), which was integrated within the *qiime deblur denoise-16S* pipeline. Rarefaction was performed at depth of 10 000 sequences per sample (at which level Shannon diversity could be robustly estimated [9]), and samples with less than 10 000 sequences were discarded. After filtering and rarefaction, a total of 54 557 ESVs in 169 samples were generated for topsoil data; a total of 44 924 ESVs in 186 samples for atlas data; and a total of 13 304 ESVs in 90 samples for farmland data. Bacterial α-diversity was represented by richness (the number of observed ESVs) and Shannon diversity. For farmland data, the whole year α-diversity was calculated based on the collective community composition where community compositions in summer and winter were merged as one community for each plot [10].

**Statistical analyses**

The linear relationships between bacterial α-diversity (richness or Shannon diversity) and temperature or precipitation seasonality were fitted by OLS regression. Pairwise associations between geographic (absolute latitude), climatic (MAT, mean annual temperature; AP, annual precipitation; TS, temperature seasonality; and PS, precipitation seasonality) variables and soil physiochemical properties (soil pH, C/N ratio; the most important two soil physiochemical factors in determining soil bacterial diversity and composition) in the two global datasets were tested using Pearson's correlation test (Fig. S1, S2, Table S1, S2). Piecewise structural equation models (SEMs) [11] were employed to address the direct and indirect effects of geographic, climatic factors and soil physiochemical properties on soil bacterial α-diversity for the two global datasets. Parameter estimation in SEMs generally requires at least 5 samples for each parameter in the model [12], therefore SEMs were not performed with farmland data due to its small sample size (n=45). Model estimation, evaluation and simplification were conducted with the '*piecewiseSEM*' package in R [11]. Shipley's test of directed separation: Fisher's $C$ statistic (if $p > 0.05$, then there are no missing associations and the model reproduces the data well) and AIC were used to evaluate the overall fit of the model. The hypothesized causational relationships were summarized in Fig. S6. Iterations of adding potentially important missing links or removing non-significant paths were repeated to obtain the parsimonious model with $p > 0.05$ for Shipley's d-separation test and the lowest AIC value for each global dataset. The details of coefficient estimation of the parsimonious models were given in Table S3-S6 for topsoil and atlas data, respectively. The significant paths with the standardized effect sizes of path coefficients and individual R square were shown in Fig. 2, Fig S4. All of the statistical analyses were conducted in R 4.0.1 [13].

13. R Core Team. R: A language and environment for statistical computing. Vienna, Austria: R Foundation for Statistical Computing; 2020

### Supplementary Figures

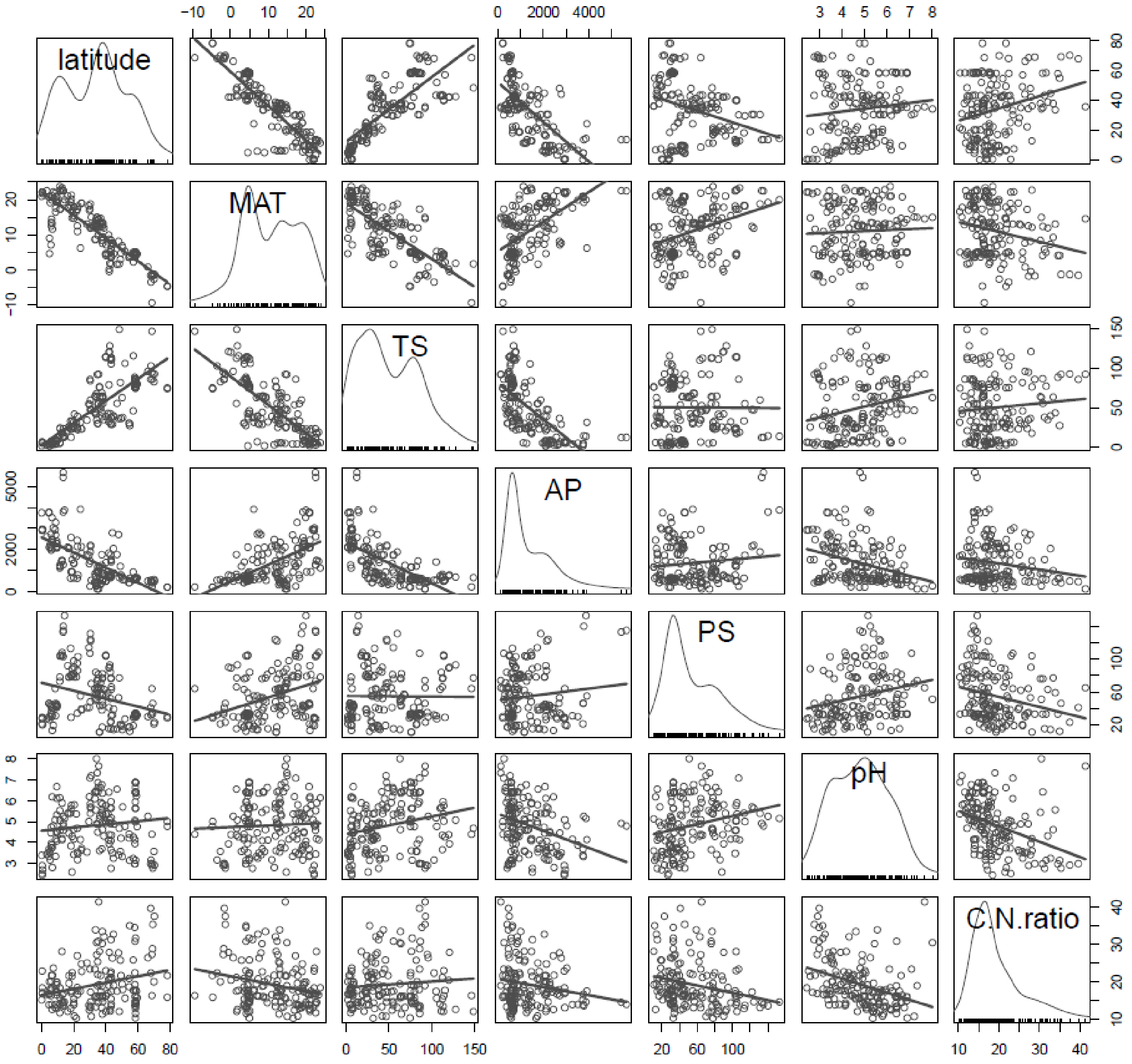


Fig. S1 Scatter plots (the upper and lower triangular panels) and adaptive kernel density estimates (the diagonal panels) for geographic (absolute latitude), climatic (MAT, mean annual temperature; TS, temperature seasonality; AP, annual precipitation; PS, precipitation seasonality) variables, and soil physiochemical properties (pH, C/N ratio) of topsoil data. Linear relationships are fitted by the OLS regression, the pairwise Pearson correlation coefficients are given in Table S1

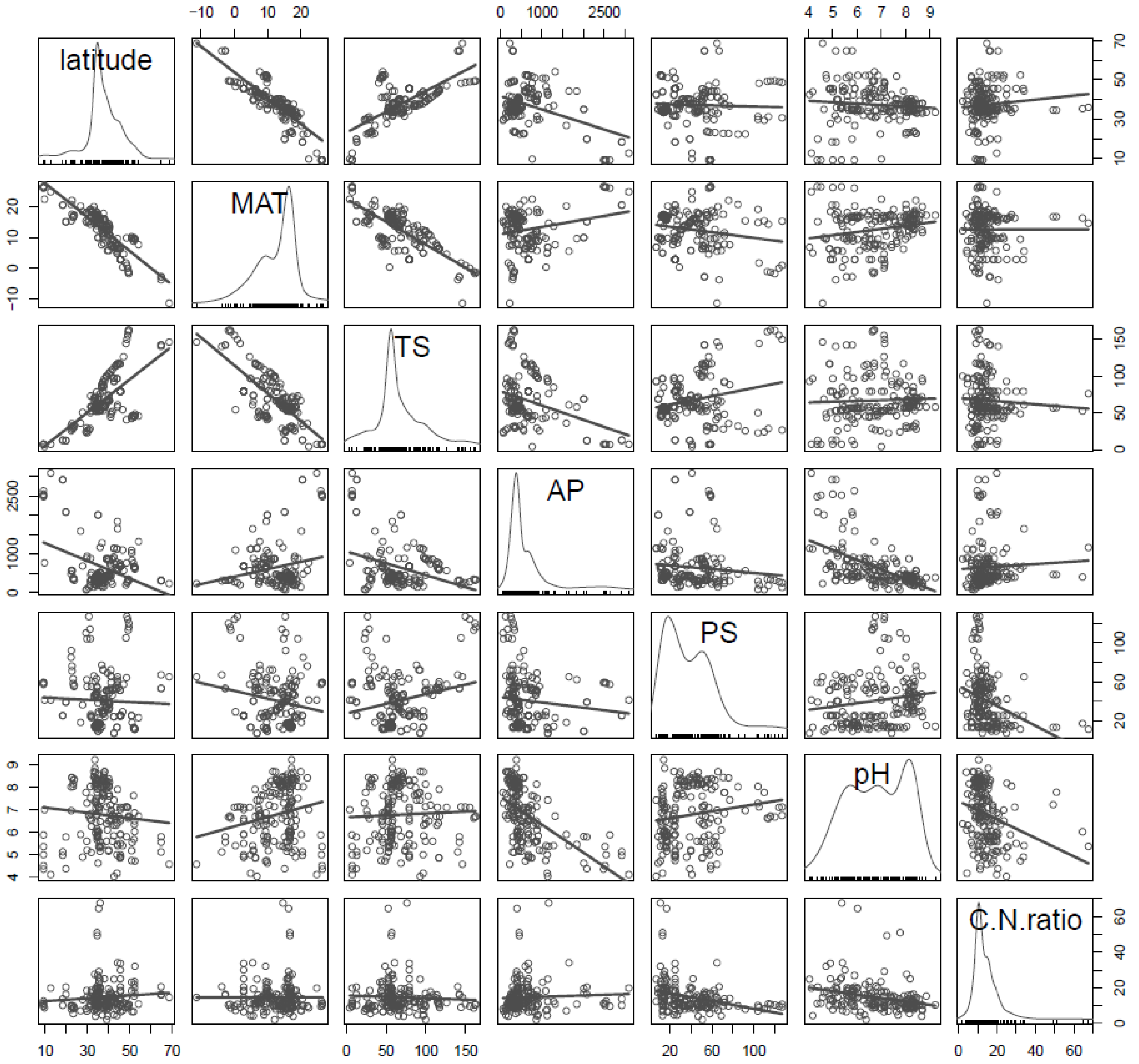


Fig. S2 Scatter plots (the upper and lower triangular panels) and adaptive kernel density estimates (the diagonal panels) for geographic (absolute latitude), climatic (MAT, mean annual temperature; TS, temperature seasonality; AP, annual precipitation; PS, precipitation seasonality) variables, and soil physiochemical properties (pH, C/N ratio) of atlas data. Linear relationships are fitted by the OLS regression, the pairwise Pearson correlation coefficients are given in Table S2

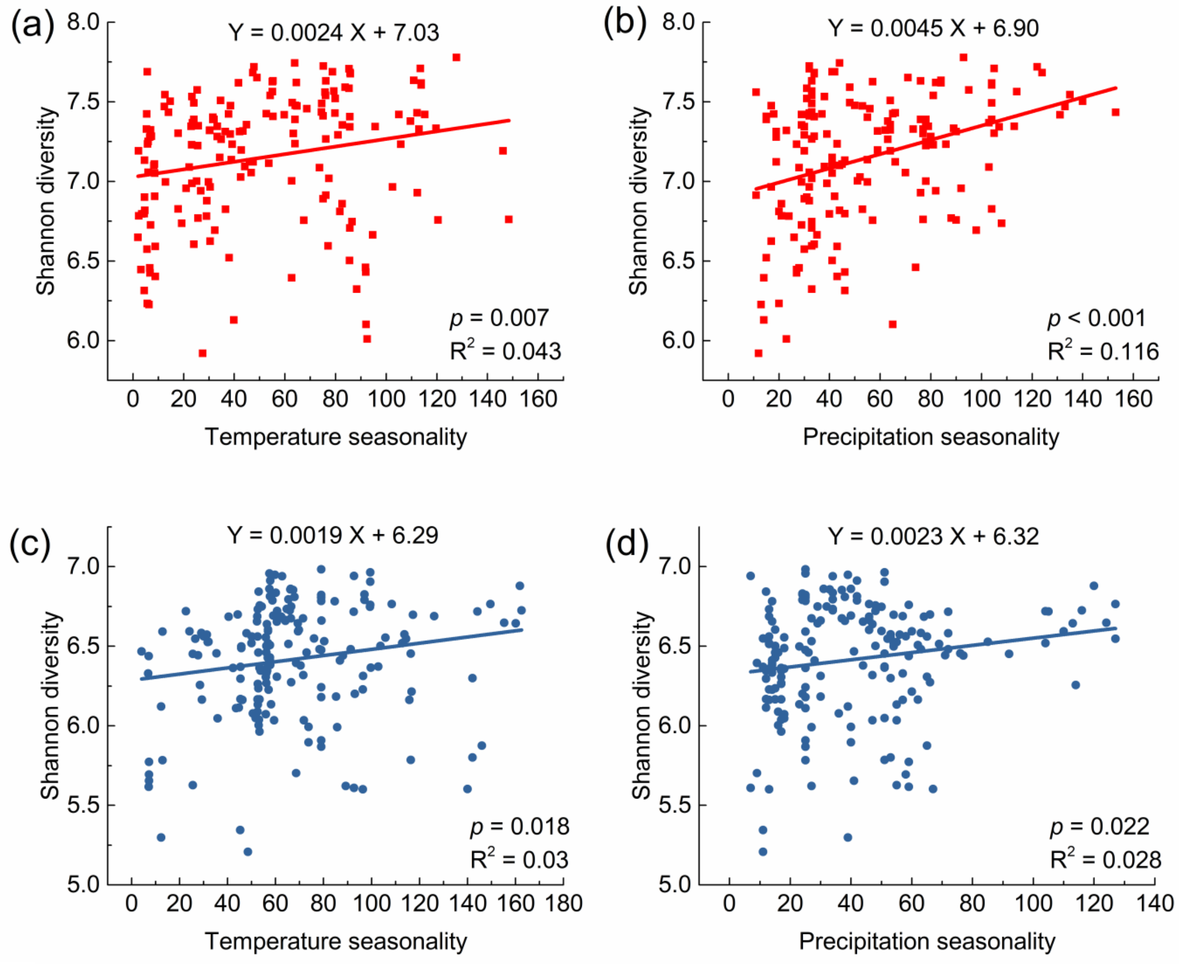


Fig. S3 The relationships between soil bacterial Shannon diversity and temperature (a, c) or precipitation (b, d) seasonality in two global datasets (topsoil data: red squares, atlas data: blue circles). Significant relationships are depicted by regression lines

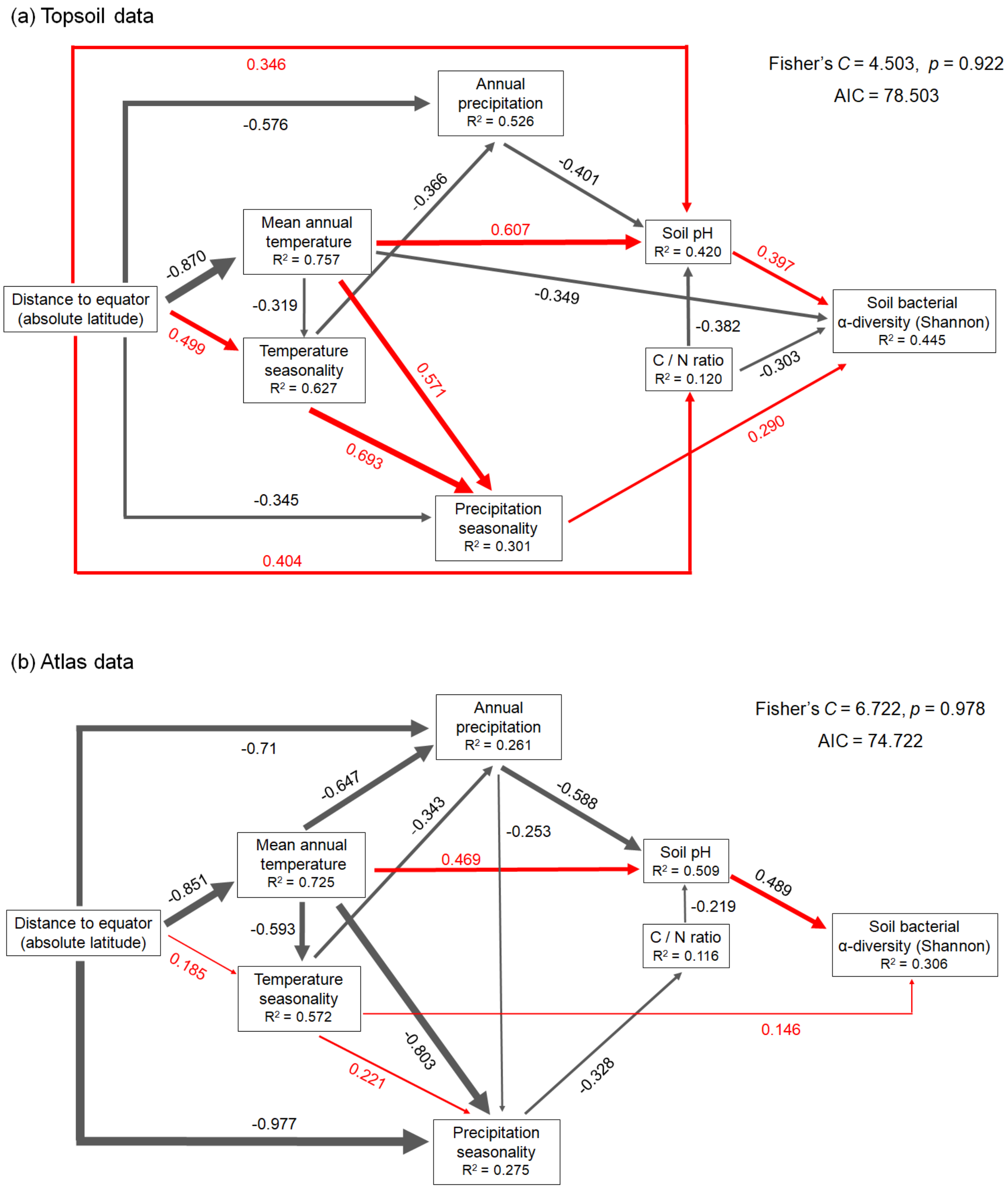


Fig. S4 Structural equation models (piecewise SEM) to address direct and indirect effects of geographic (absolute latitude), climatic (mean annual temperature, annual precipitation, temperature seasonality, precipitation seasonality) variables, and soil physiochemical properties (pH, C/N ratio) on Shannon diversity of soil bacterial communities for topsoil data (169 soil samples) (a) and atlas data (186 soil samples) (b). Red arrows represent positive paths, and black arrows represent negative paths; only significant paths are shown ($p < 0.05$). Standardized effect sizes of path coefficients are reported and indicated by path thickness. $R^2$ for component models are given in the boxes of corresponding response variables. Shipley's test of directed separation: Fisher's *C* statistic (if $p >$ 0.05, then there are no missing associations and the model reproduces the data well) and AIC are used to evaluate the overall fit of the model. Details of model evaluation, simplification and coefficients estimation are provided in supplementary methods and Table S4, S6

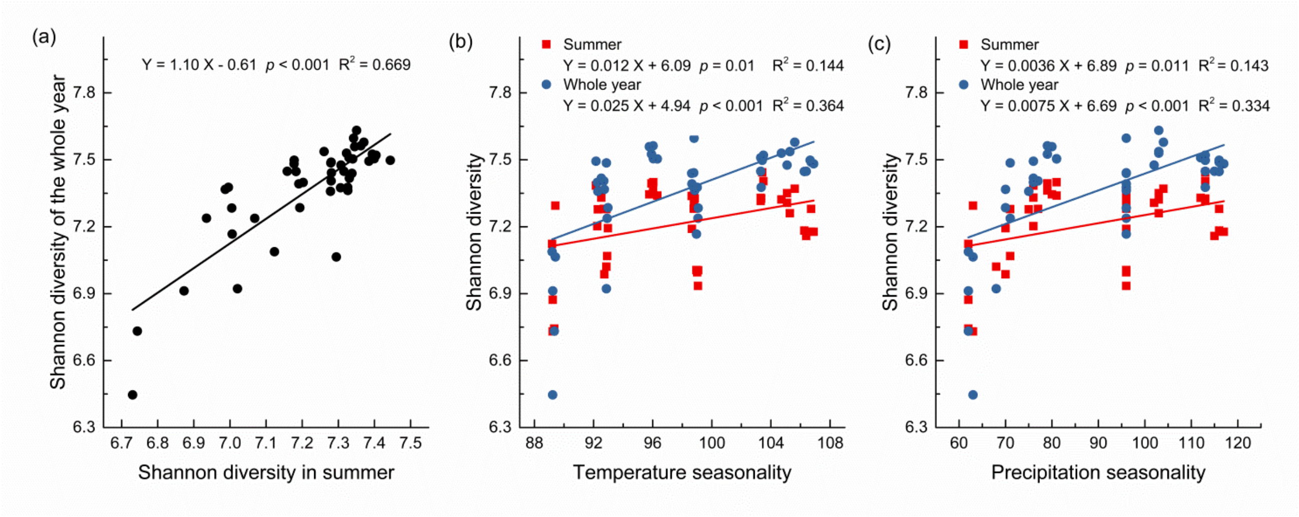


Fig. S5 The relationship between soil bacterial Shannon diversity in summer and of the whole year (a), and the relationships between soil bacterial Shannon diversity (summer: red squares; whole year: blue circles) and temperature (b) or precipitation (c) seasonality. In total of 90 soil samples were collected from 45 soil plots (geographic distances up to 873 km) across North China Plain in winter and the following summer. Shannon diversity of the whole year is calculated by merging community compositions in summer and winter for each plot. Significant relationships are depicted by regression lines

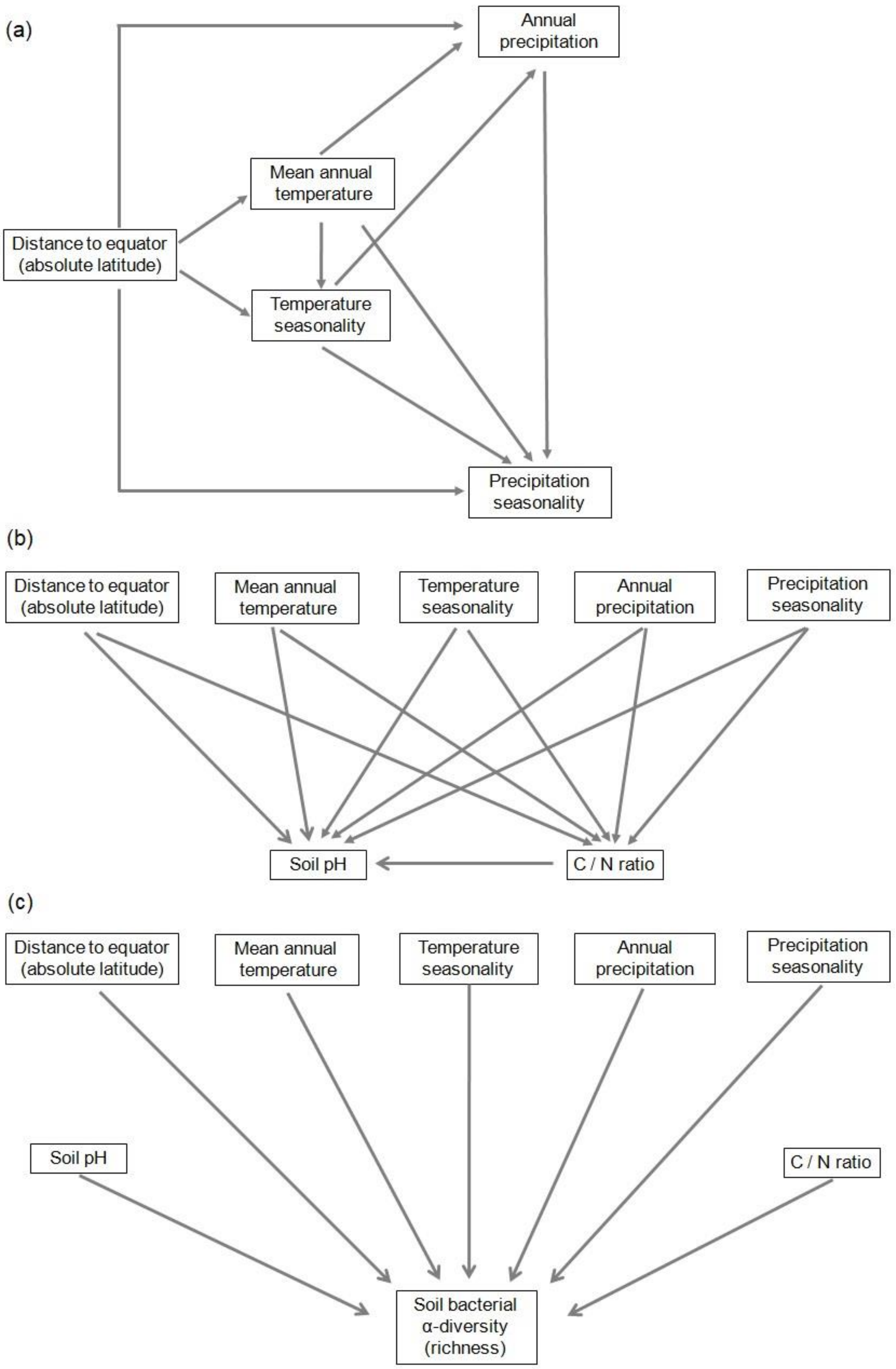


Fig. S6 A graphic illustration of the hypothesized causational relationships to be tested with piecewise structural equation model. The relationships are summarized in three sub-network: how climate factors (a), soil properties (b), and soil bacterial α-diversity (c) are affected

#### Supplementary Tables

Table S1 Pearson correlation coefficients between geographic (absolute latitude), climatic (MAT, mean annual temperature; TS, temperature seasonality; AP, annual precipitation; PS, precipitation seasonality) variables, and soil physiochemical properties (pH, C/N ratio) of topsoil data. Stars indicate significance of correlations, with * for $p < 0.05$, and ** for $p < 0.01$; $p$ values above the diagonal are adjusted for multiple tests

| | Latitude | MAT | TS | AP | PS | pH | C/N ratio |
|---|---|---|---|---|---|---|---|
| Latitude | —— | -0.87 ** | 0.78 ** | -0.69 ** | -0.31 ** | 0.12 | 0.26 * |
| MAT | -0.87 ** | —— | -0.75 ** | 0.58 ** | 0.35 ** | 0.05 | -0.24 * |
| TS | 0.78 ** | -0.75 ** | —— | -0.67 ** | -0.01 | 0.24 * | 0.09 |
| AP | -0.69 ** | 0.58 ** | -0.67 ** | —— | 0.12 | -0.34 ** | -0.18 |
| PS | -0.31 ** | 0.35 ** | -0.01 | 0.12 | —— | 0.25 * | -0.25 * |
| pH | 0.12 | 0.05 | 0.24 ** | -0.34 ** | 0.25 ** | —— | -0.37 ** |
| C/N ratio | 0.26 ** | -0.24 ** | 0.09 | -0.18* | -0.25 ** | -0.37 ** | —— |

Table S2 Pearson correlation coefficients between geographic (absolute latitude), climatic (MAT, mean annual temperature; TS, temperature seasonality; AP, annual precipitation; PS, precipitation seasonality) variables, and soil physiochemical properties (pH, C/N ratio) of atlas data. Stars indicate significance of correlations, with * for $p < 0.05$, and ** for $p < 0.01$; $p$ values above the diagonal are adjusted for multiple tests

| | Latitude | MAT | TS | AP | PS | pH | C/N ratio |
|---|---|---|---|---|---|---|---|
| Latitude | —— | -0.85 ** | 0.69 ** | -0.40 ** | -0.04 | -0.09 | 0.09 |
| MAT | -0.85 ** | —— | -0.75 ** | 0.21 * | -0.19 | 0.21 * | 0 |
| TS | 0.69 ** | -0.75 ** | —— | -0.35 ** | 0.24 * | 0.04 | -0.06 |
| AP | -0.40 ** | 0.21 ** | -0.35 ** | —— | -0.12 | -0.56 ** | 0.05 |
| PS | -0.04 | -0.19 * | 0.24 ** | -0.12 | —— | 0.16 | -0.33 ** |
| pH | -0.09 | 0.21 ** | 0.04 | -0.56 ** | 0.16 * | —— | -0.28 ** |
| C/N ratio | 0.09 | 0 | -0.06 | 0.05 | -0.33 ** | -0.28 ** | —— |

Table S3 Details of coefficient estimation of the parsimonious piecewise SEM model with geographic (absolute latitude), climatic (MAT, mean annual temperature; TS, temperature seasonality; AP, annual precipitation; PS, precipitation seasonality) variables, soil physiochemical properties (soil pH, C/N ratio) and bacterial α-diversity (richness) for topsoil data. (Fisher's $C$ = 6.588, $p$ = 0.884; AIC = 78.588). Stars indicate significance of correlations, with * for $p < 0.05$, ** for $p < 0.01$ and *** for $p < 0.001$

| Response | Predictor | Estimate | Std.Error | DF | Crit.Value | *p* value | Std.Estimate | |
|---|---|---|---|---|---|---|---|---|
| MAT | latitude | -0.3288 | 0.0145 | 166 | -22.7134 | < 0.0001 | -0.8698 | *** |
| TS | latitude | 0.8947 | 0.1728 | 165 | 5.177 | < 0.0001 | 0.4987 | *** |
| TS | MAT | -1.5139 | 0.4572 | 165 | -3.3115 | 0.0011 | -0.319 | ** |
| AP | latitude | -29.9846 | 6.1131 | 164 | -4.905 | < 0.0001 | -0.576 | *** |
| AP | MAT | -27.041 | 15.4898 | 164 | -1.7457 | 0.0827 | -0.1964 | |
| AP | TS | -10.616 | 2.5542 | 164 | -4.1564 | 0.0001 | -0.3658 | *** |
| PS | latitude | -0.5462 | 0.2263 | 164 | -2.414 | 0.0169 | -0.3445 | * |
| PS | MAT | 2.3938 | 0.5733 | 164 | 4.1752 | < 0.0001 | 0.5707 | *** |
| PS | TS | 0.612 | 0.0945 | 164 | 6.474 | < 0.0001 | 0.6925 | *** |
| C/N | TS | -0.0394 | 0.0224 | 164 | -1.758 | 0.0806 | -0.2212 | |
| C/N | PS | -0.0256 | 0.0168 | 164 | -1.5288 | 0.1282 | -0.1273 | |
| C/N | latitude | 0.1289 | 0.0422 | 164 | 3.0575 | 0.0026 | 0.4037 | ** |
| pH | MAT | 0.1006 | 0.0221 | 161 | 4.5489 | < 0.0001 | 0.607 | *** |
| pH | TS | 0.0067 | 0.0041 | 161 | 1.6585 | 0.0992 | 0.1926 | |
| pH | AP | -0.0005 | 0.0001 | 161 | -4.5899 | < 0.0001 | -0.4012 | *** |
| pH | PS | 0.0041 | 0.0029 | 161 | 1.425 | 0.1561 | 0.1029 | |
| pH | C/N | -0.0749 | 0.0126 | 161 | -5.9534 | < 0.0001 | -0.382 | *** |
| pH | latitude | 0.0217 | 0.009 | 161 | 2.4036 | 0.0174 | 0.3463 | * |
| richness | MAT | -29.3487 | 5.3437 | 163 | -5.4922 | < 0.0001 | -0.3327 | *** |
| richness | PS | 4.0998 | 1.2979 | 163 | 3.1588 | 0.0019 | 0.195 | ** |
| richness | pH | 223.4554 | 32.6104 | 163 | 6.8523 | < 0.0001 | 0.4198 | *** |
| richness | C/N | -34.2919 | 6.4697 | 163 | -5.3004 | < 0.0001 | -0.3284 | *** |

Table S4 Details of coefficient estimation of the parsimonious piecewise SEM model with geographic (absolute latitude), climatic (MAT, mean annual temperature; TS, temperature seasonality; AP, annual precipitation; PS, precipitation seasonality) variables, soil physiochemical properties (soil pH, C/N ratio) and bacterial α-diversity (Shannon) for topsoil data. (Fisher's $C$ = 4.503, $p$ = 0.922; AIC = 78.503). Stars indicate significance of correlations, with * for $p < 0.05$, ** for $p < 0.01$ and *** for $p < 0.001$

| Response | Predictor | Estimate | Std.Error | DF | Crit.Value | *p* value | Std.Estimate | |
|---|---|---|---|---|---|---|---|---|
| MAT | latitude | -0.3288 | 0.0145 | 166 | -22.7134 | < 0.0001 | -0.8698 | *** |
| TS | latitude | 0.8947 | 0.1728 | 165 | 5.177 | < 0.0001 | 0.4987 | *** |
| TS | MAT | -1.5139 | 0.4572 | 165 | -3.3115 | 0.0011 | -0.319 | ** |
| AP | latitude | -29.9846 | 6.1131 | 164 | -4.905 | < 0.0001 | -0.576 | *** |
| AP | MAT | -27.041 | 15.4898 | 164 | -1.7457 | 0.0827 | -0.1964 | |
| AP | TS | -10.616 | 2.5542 | 164 | -4.1564 | 0.0001 | -0.3658 | *** |
| PS | latitude | -0.5462 | 0.2263 | 164 | -2.414 | 0.0169 | -0.3445 | * |
| PS | MAT | 2.3938 | 0.5733 | 164 | 4.1752 | < 0.0001 | 0.5707 | *** |
| PS | TS | 0.612 | 0.0945 | 164 | 6.474 | < 0.0001 | 0.6925 | *** |
| C/N | TS | -0.0394 | 0.0224 | 164 | -1.758 | 0.0806 | -0.2212 | |
| C/N | PS | -0.0256 | 0.0168 | 164 | -1.5288 | 0.1282 | -0.1273 | |
| C/N | latitude | 0.1289 | 0.0422 | 164 | 3.0575 | 0.0026 | 0.4037 | ** |
| pH | MAT | 0.1006 | 0.0221 | 161 | 4.5489 | < 0.0001 | 0.607 | *** |
| pH | TS | 0.0067 | 0.0041 | 161 | 1.6585 | 0.0992 | 0.1926 | |
| pH | AP | -0.0005 | 0.0001 | 161 | -4.5899 | < 0.0001 | -0.4012 | *** |
| pH | PS | 0.0041 | 0.0029 | 161 | 1.425 | 0.1561 | 0.1029 | |
| pH | C/N | -0.0749 | 0.0126 | 161 | -5.9534 | < 0.0001 | -0.382 | *** |
| pH | latitude | 0.0217 | 0.009 | 161 | 2.4036 | 0.0174 | 0.3463 | * |
| Shannon | MAT | -0.0191 | 0.006 | 162 | -3.2059 | 0.0016 | -0.3491 | ** |
| Shannon | TS | -0.0014 | 0.0012 | 162 | -1.1814 | 0.2392 | -0.1242 | |
| Shannon | PS | 0.0038 | 0.0009 | 162 | 4.1737 | < 0.0001 | 0.2902 | *** |
| Shannon | pH | 0.1314 | 0.0228 | 162 | 5.7579 | < 0.0001 | 0.3972 | *** |
| Shannon | C/N | -0.0197 | 0.0042 | 162 | -4.6418 | < 0.0001 | -0.303 | *** |

Table S5 Details of coefficient estimation of the parsimonious piecewise SEM model with geographic (absolute latitude), climatic (MAT, mean annual temperature; TS, temperature seasonality; AP, annual precipitation; PS, precipitation seasonality) variables, soil physiochemical properties (soil pH, C/N ratio) and bacterial α-diversity (richness) for atlas data. (Fisher's $C = 9.032$, $p = 0.912$; AIC = 77.032). Stars indicate significance of correlations, with * for $p < 0.05$ and *** for $p < 0.001$

| Response | Predictor | Estimate | Std.Error | DF | Crit.Value | *p* value | Std.Estimate | |
|---|---|---|---|---|---|---|---|---|
| MAT | latitude | -0.5509 | 0.025 | 184 | -22.006 | < 0.0001 | -0.8513 | *** |
| TS | latitude | 0.6025 | 0.3006 | 183 | 2.0042 | 0.0465 | 0.1846 | * |
| TS | MAT | -2.991 | 0.4645 | 183 | -6.4388 | < 0.0001 | -0.5932 | *** |
| AP | latitude | -41.0734 | 7.1014 | 182 | -5.7838 | < 0.0001 | -0.71 | *** |
| AP | MAT | -57.8445 | 12.0227 | 182 | -4.8113 | < 0.0001 | -0.647 | *** |
| AP | TS | -6.082 | 1.7275 | 182 | -3.5207 | 0.0005 | -0.343 | *** |
| PS | latitude | -2.6587 | 0.3608 | 181 | -7.3696 | < 0.0001 | -0.9774 | *** |
| PS | MAT | -3.3752 | 0.596 | 181 | -5.6632 | < 0.0001 | -0.8029 | *** |
| PS | TS | 0.1843 | 0.0834 | 181 | 2.2113 | 0.0283 | 0.2211 | * |
| PS | AP | -0.0119 | 0.0035 | 181 | -3.4352 | 0.0007 | -0.2528 | *** |
| C/N | latitude | 0.0677 | 0.0613 | 183 | 1.1042 | 0.271 | 0.0768 | |
| C/N | PS | -0.1064 | 0.0225 | 183 | -4.721 | < 0.0001 | -0.3284 | *** |
| pH | MAT | 0.0927 | 0.0157 | 180 | 5.896 | < 0.0001 | 0.4686 | *** |
| pH | TS | 0.0063 | 0.0033 | 180 | 1.9206 | 0.0564 | 0.1599 | |
| pH | AP | -0.0013 | 0.0001 | 180 | -10.521 | < 0.0001 | -0.5883 | *** |
| pH | PS | 0.0034 | 0.0027 | 180 | 1.2782 | 0.2028 | 0.0729 | |
| pH | C/N | -0.0318 | 0.0081 | 180 | -3.938 | 0.0001 | -0.2188 | *** |
| richness | TS | 2.558 | 0.7296 | 182 | 3.5061 | 0.0006 | 0.191 | *** |
| richness | pH | 201.2829 | 19.341 | 182 | 10.4071 | < 0.0001 | 0.5898 | *** |
| richness | C/N | -6.8648 | 2.8112 | 182 | -2.442 | 0.0156 | -0.1385 | * |

Table S6 Details of coefficient estimation of the parsimonious piecewise SEM model with geographic (absolute latitude), climatic (MAT, mean annual temperature; TS, temperature seasonality; AP, annual precipitation; PS, precipitation seasonality) variables, soil physiochemical properties (soil pH, C/N ratio) and bacterial α-diversity (Shannon) for atlas data. (Fisher's $C = 6.722$, $p = 0.978$; AIC = 74.722). Stars indicate significance of correlations, with * for $p < 0.05$ and *** for $p < 0.001$

| Response | Predictor | Estimate | Std.Error | DF | Crit.Value | *p* value | Std.Estimate | |
|---|---|---|---|---|---|---|---|---|
| MAT | latitude | -0.5509 | 0.025 | 184 | -22.006 | < 0.0001 | -0.8513 | *** |
| TS | latitude | 0.6025 | 0.3006 | 183 | 2.0042 | 0.0465 | 0.1846 | * |
| TS | MAT | -2.991 | 0.4645 | 183 | -6.4388 | < 0.0001 | -0.5932 | *** |
| AP | latitude | -41.0734 | 7.1014 | 182 | -5.7838 | < 0.0001 | -0.71 | *** |
| AP | MAT | -57.8445 | 12.0227 | 182 | -4.8113 | < 0.0001 | -0.647 | *** |
| AP | TS | -6.082 | 1.7275 | 182 | -3.5207 | 0.0005 | -0.343 | *** |
| PS | latitude | -2.6587 | 0.3608 | 181 | -7.3696 | < 0.0001 | -0.9774 | *** |
| PS | MAT | -3.3752 | 0.596 | 181 | -5.6632 | < 0.0001 | -0.8029 | *** |
| PS | TS | 0.1843 | 0.0834 | 181 | 2.2113 | 0.0283 | 0.2211 | * |
| PS | AP | -0.0119 | 0.0035 | 181 | -3.4352 | 0.0007 | -0.2528 | *** |
| C/N | latitude | 0.0677 | 0.0613 | 183 | 1.1042 | 0.271 | 0.0768 | |
| C/N | PS | -0.1064 | 0.0225 | 183 | -4.721 | < 0.0001 | -0.3284 | *** |
| pH | MAT | 0.0927 | 0.0157 | 180 | 5.896 | < 0.0001 | 0.4686 | *** |
| pH | TS | 0.0063 | 0.0033 | 180 | 1.9206 | 0.0564 | 0.1599 | |
| pH | AP | -0.0013 | 0.0001 | 180 | -10.521 | < 0.0001 | -0.5883 | *** |
| pH | PS | 0.0034 | 0.0027 | 180 | 1.2782 | 0.2028 | 0.0729 | |
| pH | C/N | -0.0318 | 0.0081 | 180 | -3.938 | 0.0001 | -0.2188 | *** |
| Shannon | TS | 0.0016 | 0.0007 | 182 | 2.3658 | 0.019 | 0.1464 | * |
| Shannon | pH | 0.1397 | 0.0184 | 182 | 7.5923 | < 0.0001 | 0.4887 | *** |
| Shannon | C/N | -0.0042 | 0.0027 | 182 | -1.5712 | 0.1179 | -0.1012 | |
